# ErrorX: automated error correction for immune repertoire sequencing datasets

**DOI:** 10.1101/2020.02.17.952408

**Authors:** Alexander M Sevy

## Abstract

**Motivation:** Recent advances in DNA sequencing technology have allowed deep profiling of B- and T-cell receptor sequences on an unprecedented scale. However, sequencing errors pose a significant challenge in expanding the scope of these experiments. Errors can arise both by PCR during library preparation and by miscalled bases on the sequencing instrument itself. These errors compromise the validity of biological conclusions drawn from the data.

**Results:** To address these concerns I have developed ErrorX, a software for automated error correction of B- and T-cell receptor NGS datasets. ErrorX uses deep learning to automatically identify bases that have a high probability of being erroneous. In benchmark studies, ErrorX reduced the overall error rate of public datasets by up to 36% with a false positive rate of 0.05% or less. Since ErrorX is a pure bioinformatics approach, it can be directly applied to any existing antibody or T-cell receptor sequencing datasets to infer sites of probable error without any changes in library preparation.

**Availability:** ErrorX is free for non-commercial use, with both a command-line interface and GUI available for Mac, Linux, and Windows operating systems, and full documentation available. Pre-compiled binaries are available at https://endeavorbio.com/downloads/.

## Introduction

The scale of immune repertoire sequencing (RepSeq) has increased exponentially over the past decade. Sequencing platforms such as the Illumina MiSeq, HiSeq, and NovaSeq are able to generate over 10^9^ sequencing reads per sample (Briney *et al.*, 2019; Soto *et al.*, 2019). There are currently over 4.0 terabases of publicly available B- and T-cell receptor (BCR and TCR, respectively) sequences available in the NCBI Sequence Read Archive (SRA; Leinonen *et al.*, 2011) and data continues to be added every year. In addition, initiatives such as the Human Vaccines Project rely on unprecedented deep sequencing of human donors to decode the human immune system (Soto *et al.*, 2019; Wooden and Koff, 2018).

However, with the promise of large-scale immune repertoire sequencing come challenges as well. In particular, sequencing error is a fundamental technical limitation that limits the scale and scope of RepSeq experiments. Sequencing error can arise both from PCR errors during library preparation and miscalled bases on the sequencing instrument. Illumina reports an error rate of roughly 0.1% for MiSeq and HiSeq instruments, *i.e.* 1 error per kb sequenced (Glenn, 2011). This is compounded by the error rate of the DNA polymerase used during library preparation, which can vary from 10^−5^ to 10^−6^ errors per base per cycle, even when using a high-fidelity polymerase (Hestand *et al.*, 2016; McInerney *et al.*, 2014). Spike-in studies have shown that 5% of high-quality sequences in RepSeq datasets contain erroneous reads, which can lead to over 100 erroneous variants per clone (Shugay *et al.*, 2014; Friedensohn *et al.*, 2018).

Errors in antibody sequencing datasets are especially problematic because errors are indistinguishable from somatic hypermutation. Somatic variants provide a great deal of biological insight, since they have been mutated in response to a challenge and can provide information on the specificity of clones. This creates a difficult situation where biologically interesting somatic variants are confounded with error-ridden sequences. It has been shown in several cases that elaborate clonal lineages can be created from a single DNA template by error alone (Briney *et al.*, 2016; Khan *et al.*, 2016).

The predominant approach to error correction in RepSeq datasets uses barcoding via unique molecular identifiers (UMIs) attached to the DNA strand during library preparation to identify reads that have originated from the same template. Reads with matching UMIs can then be collapsed to a consensus sequence to eliminate errors. UMIs have been implemented in many varieties in the RepSeq community with success (Shugay *et al.*, 2014; Friedensohn *et al.*, 2018; Briney *et al.*, 2016; Khan *et al.*, 2016; Turchaninova *et al.*, 2016; Cole *et al.*, 2016). However, this technology still faces several limitations. UMI collisions, where two distinct templates contain the same barcode, can be common depending on the design of the barcode and must be accounted for by complex bioinformatic pipelines. In addition, consensus sequences cannot be generated with confidence for barcodes with less than five reads, potentially eliminating a significant portion of the dataset. In addition, incorporating UMIs into a library preparation protocol can be technically challenging, leading to complex workflows. Lastly, since UMIs must be incorporated before sequencing, they do not address the problem of error in legacy datasets.

Existing bioinformatic approaches to error correction use abundance-based clustering to filter out rare clones and keep those that are represented by multiple reads. These approaches have been most successful for correction of T-cell receptors, which are more redundant due to the lack of somatic hypermutation (Gerritsen *et al.*, 2016; Giraud *et al.*, 2014; Bolotin *et al.*, 2015). However, these approaches are limited in that they discard rare clones, which may be of significant interest in antibody sequencing datasets. This can result in a tremendous loss of data. Although there are many software tools for reconstruction and analysis of BCRs (Briney *et al.*, 2016; Vander Heiden *et al.*, 2014; Safonova *et al.*, 2015; Kovaltsuk *et al.*, 2018), there is currently no option for error correction of unlabeled BCR sequences.

To address these concerns, I have developed ErrorX, an automated suite for immune repertoire error correction. ErrorX uses deep learning to identify individual nucleotide positions that are likely to have arisen from either PCR or sequencing error. Nucleotides with a high probability of being erroneous are annotated and can be accounted for when building clonal lineages, estimating total diversity, or measuring degree of somatic hypermutation. ErrorX does not use any clustering or consensus-building to assign errors, which confers several advantages over existing approaches: 1) rare clones are maintained in the datasets, 2) high sequencing depth is not necessary for error correction with confidence, and 3) processing time scales linearly with number of sequences, allowing for rapid error correction even in large datasets. In addition, ErrorX is trained on real datasets, not synthetic repertoires, meaning that the results shown in benchmarking translate directly to the performance on a new dataset. Since ErrorX is a pure bioinformatics approach to error correction, it does not rely on UMI barcodes and operates directly on unlabeled sequences, making it amenable to correction of legacy data. To the author’s knowledge, this is the first method for error correction of unlabeled BCR repertoires.

## Results

### Error correction workflow

The ErrorX workflow is summarized in Figure 1. Annotated BCR sequences from public datasets were used to train a deep neural network to predict the probability that a nucleotide at a given position is erroneous. Features such as nucleotide identity, Phred quality score (a quality metric reported by the sequencing instrument), and germline nucleotide identity were calculated from a sliding window surrounding the position of interest and used as input to the neural network (see Methods for details). Since the training datasets were fully annotated, the problem was formulated as a supervised binary classification task of predicting a given base as error/no error, and performance was measured on an independent validation dataset. Once the network was optimized to maximize performance, the same network was then applied to unlabeled data to identify erroneous nucleotides.

**Figure 1.**
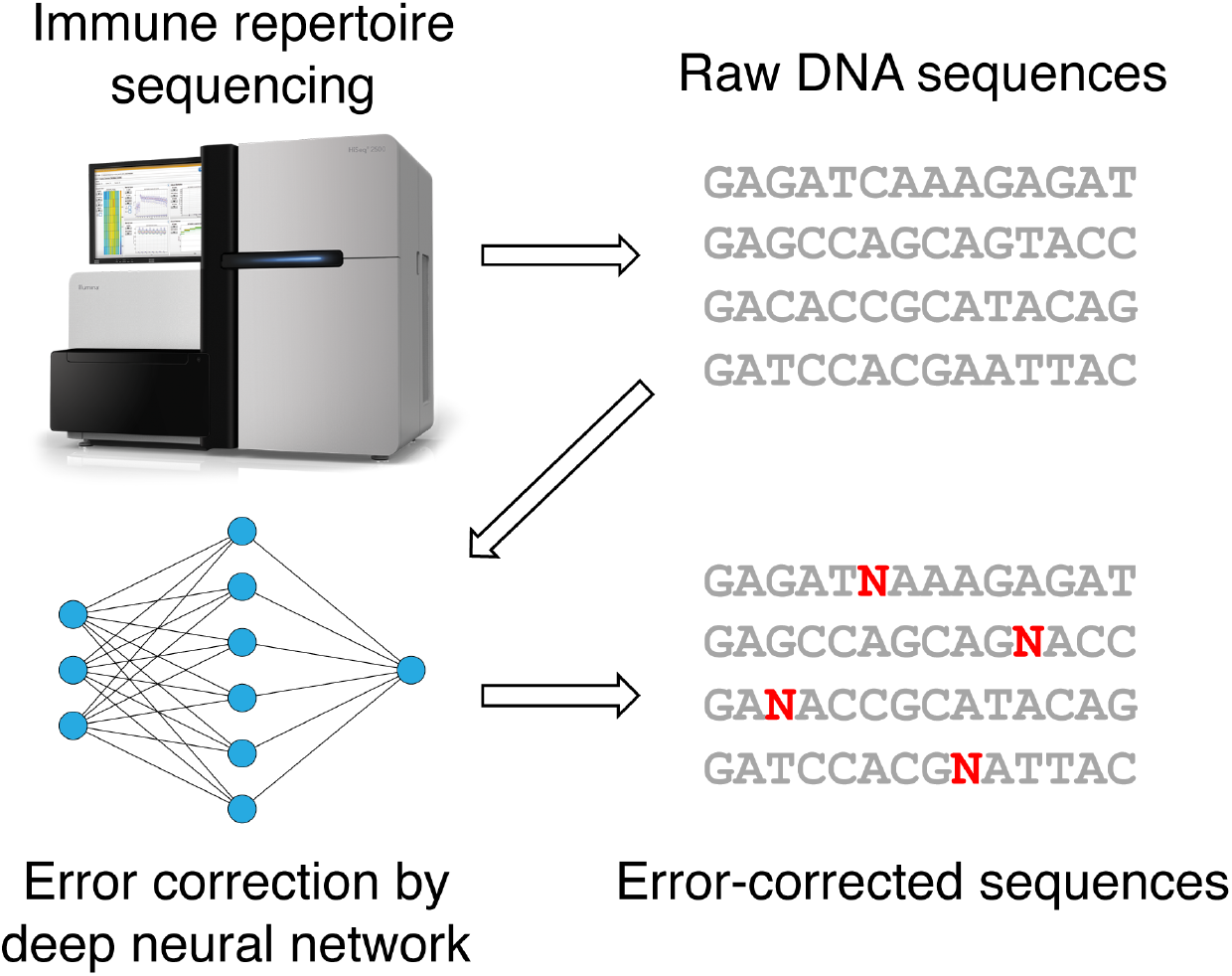
ErrorX error correction workflow. Raw sequence data of antibodies or T-cell receptors is given as input to a deep neural network trained to predict sites of sequencing or PCR error. The input sequences are then annotated with positions where error is likely for the user to interpret in downstream experiments.

### Performance on public datasets

After training the ErrorX neural network on human and mouse BCR sequences, the performance was measured on validation datasets of human and mouse BCR sequences from donors not included in the training set. In addition, ErrorX was tested on a validation dataset of human TCR sequences. It is worth noting that ErrorX was never explicitly trained on TCR sequence data - if the rules defining probable sites of error are universal across receptor type, then performance should be equally robust for BCR and TCR data. ErrorX performed very well in discriminating between errors and non-errors in all three datasets (Figure 2a), with area under the curve on the receiver operating curve (ROC AUC) ranging from 0.92 - 0.97. Importantly, the false positive rates at the optimized threshold were 0.05% or less in all cases, meaning that while error positions were accurately identified, it was rare that a correct base was called an error. As a control, I plotted the performance of Phred quality scores alone as a discriminator. ErrorX significantly outperformed the naive approach of using quality scores alone with a ROC AUC of 0.6.

**Figure 2.**
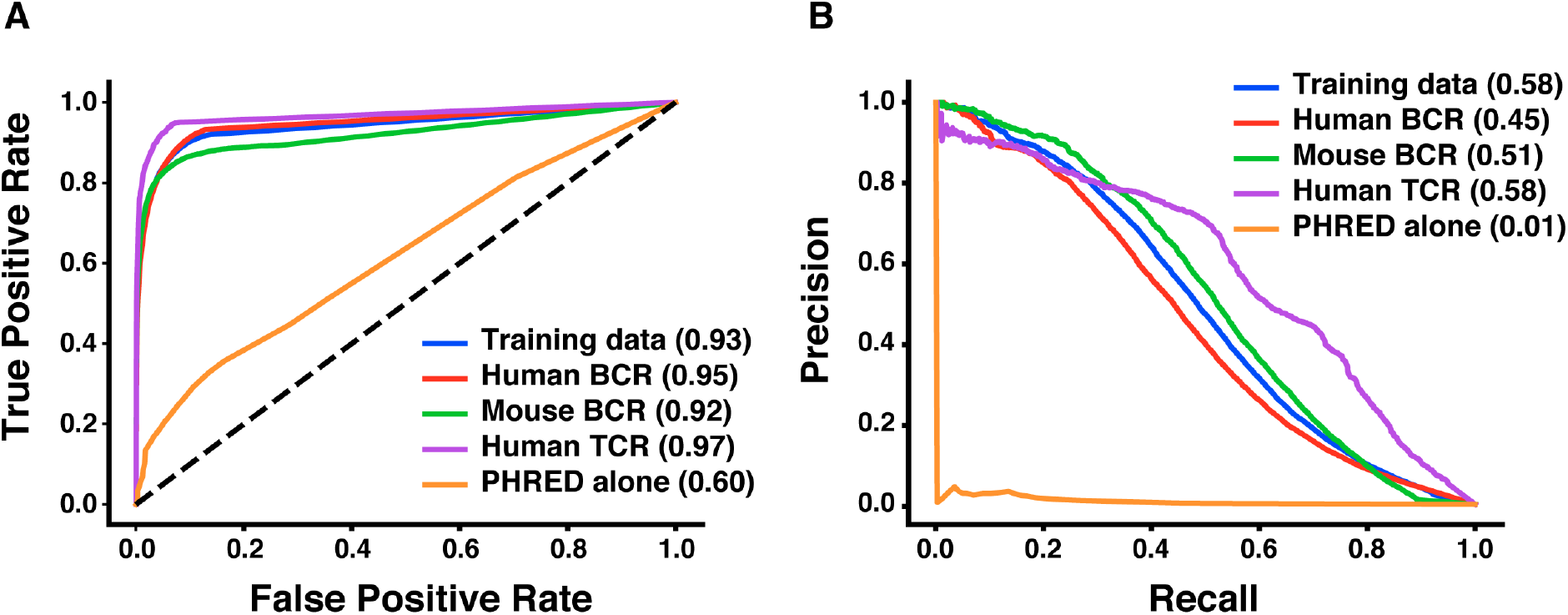
Performance of the ErrorX network in error correction. The ErrorX neural network was trained on a set of human and mouse BCR sequences (labeled “Training data”) and evaluated on three validation datasets, of either human BCRs, mouse BCRs, or human TCRs. A. ROC curve showing performance in error prediction for the four datasets. Phred quality score alone was used as a naïve control, as it was the most predictive input feature. ROC area under the curve (AUC) is shown in parentheses. B. Precision-recall curves for error prediction. Average precision score is shown in parentheses.

Error identification is a highly class-imbalanced problem, with many more negative data points than positives (*i.e.* many more correct bases than errors). As ROC AUC can be misleading in class-imbalanced problems, I also plotted the precision-recall (PR) curves for these datasets and calculated the average precision score (Figure 2b, average precision score shown in parenthesis). In agreement with the ROC curves, performance in precision-recall space was very robust, with average precision score ranging from 0.45 - 0.58. Based on these precision-recall curves, an optimized threshold was calculated where precision in the validation set was at least 0.75 - this is the probability cutoff used throughout the ErrorX protocol. At the optimized threshold ErrorX achieved precision ranging from 75 - 85% with recall ranging from 29 - 36% (Table 1). As in the ROC curve, quality scores alone did not provide robust discrimination in PR space.

**Table 1.**
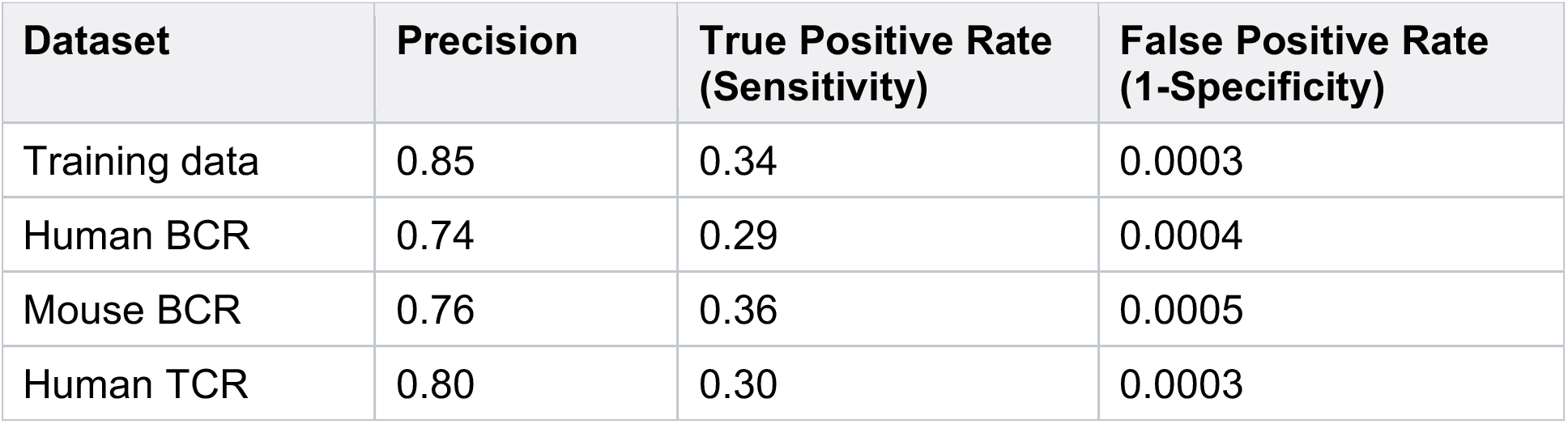
Performance of the ErrorX network at the optimized probability cutoff against the training dataset and three validation datasets.

### Neural network architecture

After parameter optimization, the final network consisted of 3 hidden layers of 256, 128, and 64 nodes. To justify the use of a deep neural network, and verify that a simpler model could not reproduce the same results, I trained additional networks consisting of either 1) a single hidden layer with 256 nodes, or 2) no hidden layer (logistic regression model). The optimized deep network outperformed both of these alternative models, both in ROC AUC and average precision scores (Table 2). This indicates that error correction is indeed a complex problem with non-linear relationships between the input features, and greatly benefits from the use of a deep neural network.

**Table 2:**
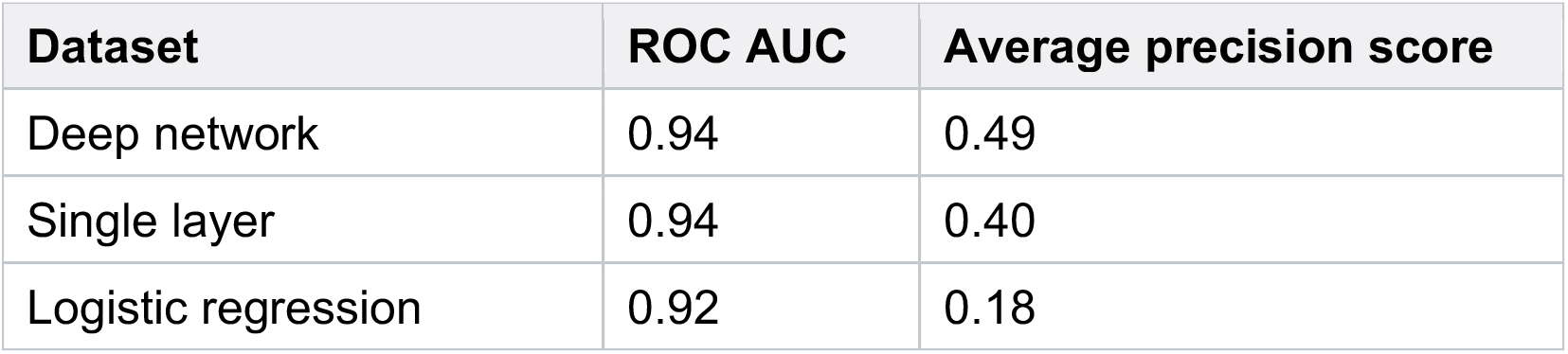
Performance of the optimized, deep neural network vs. simpler models. The deep network consisted of 3 hidden layers of 256, 128, and 64 nodes, whereas the single layer model had a single hidden layer of 256 nodes, and the logistic regression model had no hidden layer.

| Dataset | ROC AUC | Average precision score |
| --- | --- | --- |
| Deep network | 0.94 | 0.49 |
| Single layer | 0.94 | 0.40 |
| Logistic regression | 0.92 | 0.18 |

### Computational performance

Since ErrorX does not cluster or group sequences in any way before error correction, the runtime required to process sequences scales linearly with the number of sequences. In benchmark studies, 1,000 sequences could be error-corrected in 18.5 seconds when run from the raw FASTQ file (Table 3). The majority of the runtime is spent on germline gene assignment by IgBlast and not in actual error correction. Therefore the pipeline can be sped up significantly when the germline has been pre-assigned and the data input to ErrorX as a TSV file – the runtime in this case was 3.2 seconds (Table 3). Based on this computational performance, ErrorX can easily be integrated into any processing pipeline, with minimal effect on total runtime.

**Table 3:**
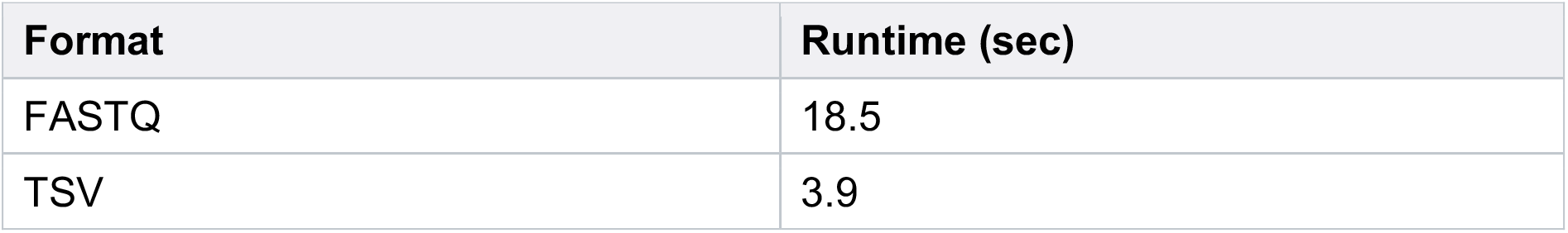
Runtime required for processing 1,000 sequences with the ErrorX protocol, based on using either raw FASTQ data as input, or a TSV file with pre-annotated sequences where the germline had already been assigned.

### False clonal lineage reduction

A frequent problem in immune repertoire sequencing dataset is the identification of spurious clonal lineages that are caused not by somatic hypermutation, but errors introduced during the sequencing process. The presence of such erroneous lineages has been illustrated in multiple cases (Briney *et al.*, 2016; Khan *et al.*, 2016) and is currently a major obstacle to clonal lineage analysis in non-error corrected data. As an illustration of the power of ErrorX in identifying erroneous sequences, I applied ErrorX to a spurious clonal lineage in a mouse BCR sequencing dataset from the independent validation set. This lineage consisted of eight “clones” which were distinct on the nucleotide level, but were known to have originated from the same DNA template based on a UMI barcode (Figure 3a). When ErrorX was applied to these spurious clones, the neural network correctly identified the positions of error in all eight clones, without misidentifying any correct bases as errors (Figure 3b). Importantly, this error correction was done without using the UMI barcode as an input, purely based on the unlabeled nucleotide sequence. In this case, the spurious clonal lineage of eight clones could be accurately characterized as erroneous by ErrorX and reduced to the one correct sequence. This shows that ErrorX is an important tool to incorporate in any datasets where clonal lineages are being generated, especially when UMI barcodes are not present, to eliminate the possibility of analyzing false clonal lineages.

**Figure 3.**
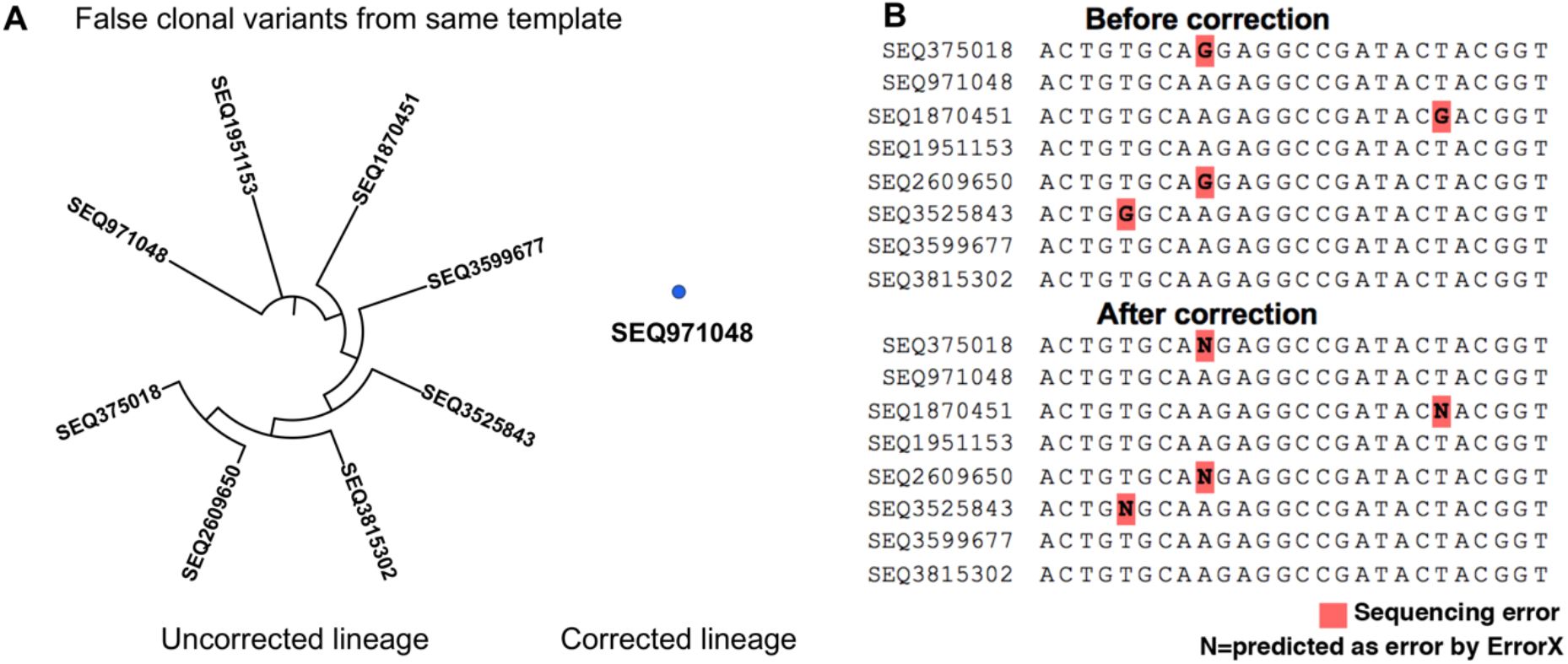
Case study of ErrorX used to correct spurious clonal lineages. A. A family of eight sequences with a common UMI barcode, but different nucleotide sequences resulting from sequencing or PCR error, was identified. After UMI removal and error correction by ErrorX, the erroneous bases were correctly annotated and the spurious clonal lineage was reduced to a single member (SEQ971048). B. Illustration of error correction in sequences from panel A. Nucleotides known to be erroneous by UMI-based consensus are highlighted in red. N nucleotides indicate those predicted to be errors by ErrorX. In this case study, all sequencing errors were correctly detected by ErrorX, with no false positives (*i.e.* correct nucleotides labeled as erroneous).

### GUI integration

In addition to a command line interface, ErrorX is also available as a graphical user interface (GUI), called ErrorX Viewer. All of the functionality from the command line is also available in the GUI, including error correction, V, D, and J gene assignment, CDR1, 2, and 3 length analyses, and clonotype analysis. In addition to error correction, ErrorX Viewer provides a full suite of sequence analysis for any stage of antibody discovery or repertoire sequencing. ErrorX Viewer has a tab layout showing 1) a summary of the input data, including number of total sequences, unique sequences, and productive sequences, 2) a summary of IGHV and IGHJ gene usage, 3) histograms portraying the lengths of CDR1, CDR2, and CDR3 loops, 4) a summary of the error profile of the dataset in addition to estimates of the total error rate, 5) dominant clonotypes contained in the dataset and their relative proportions, and 6) a full report of IgBlast output for the input dataset. A screenshot of the ErrorX Viewer interface, showing the error rate analysis tab, is shown in Figure 4. The analysis can be run over multiple threads on the host computer, allowing for rapid processing time.

**Figure 4.**
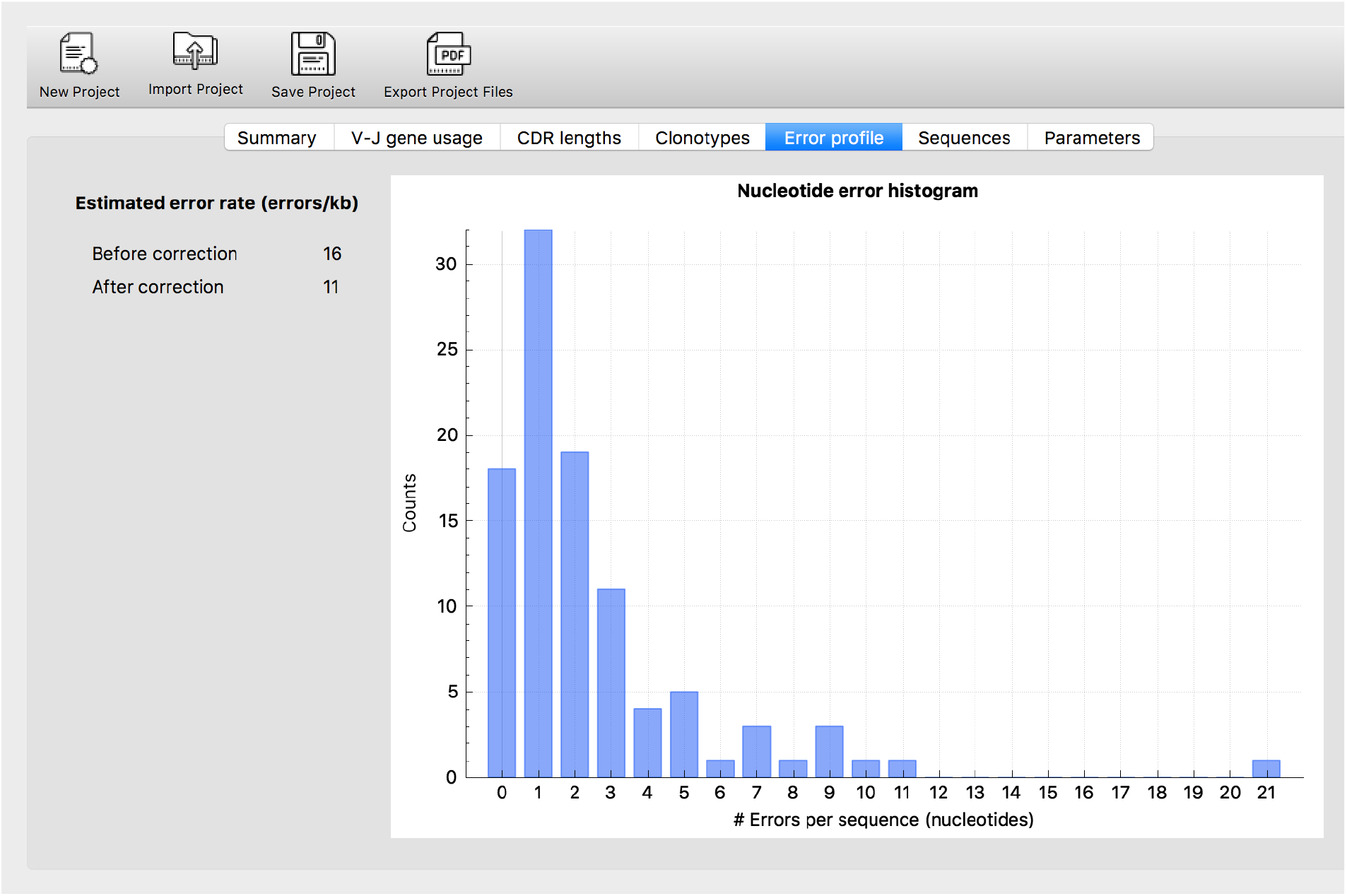
Error profile window from the ErrorX Viewer GUI. After error correction a profile of sequencing errors in the dataset is shown in the ErrorX Viewer GUI window. Shown is a histogram of number of predicted nucleotide errors per sequence. In addition the total error rate in the dataset is estimated in terms of number of errors per kilobase (left panel), as well as the estimated error rate after correction.

### Data types supported

Currently ErrorX supports next-generation sequencing data of BCRs and TCRs, gathered on Illumina instruments, of human and mouse origin. ErrorX can analyze antibody heavy and light chains and TCR alpha and beta chains. ErrorX can operate directly on FASTQ data, performing germline assignment using the IgBlast algorithm (Ye *et al.*, 2013). However, if the germline has previously been assigned, a tab-separated value (TSV) file can be used as input, consisting of the following four columns: 1) sequence ID, 2) nucleotide sequence, 3) germline nucleotide sequence, and 4) Phred quality scores. Since germline assignment is the most time-consuming step of the ErrorX protocol, bypassing this step is an easy way to integrate ErrorX into an existing pipeline. ErrorX can also be applied to FASTA data - however, since Phred quality scores are an important input feature, error correction cannot be performed with FASTA input, and only the germline assignment and annotation functionality is available.

## Discussion

In this report I describe the ErrorX software, which uses deep learning to identify erroneous nucleotides in RepSeq datasets consisting of either BCR or TCR sequences. ErrorX was able to reduce the error rate by up to 36% in public datasets with a false positive rate of 0.05%. In a case study, applying ErrorX to an erroneous clonal lineage from a BCR sequencing dataset was able to correctly identify that 8 false clonal variants originated from the same parent clone, using a pure bioinformatics approach. ErrorX operates on unlabeled sequences directly from the FASTQ sequence file, allowing it to be easily integrated into an existing workflow.

## Limitations

Currently ErrorX only supports data gathered on Illumina instruments. Since the neural network was trained exclusively on Illumina sequence data, the features that contribute to erroneous positions are likely to be specific to Illumina sequencing chemistry. Sequencing data gathered on other instruments with different sequencing chemistries therefore would not be compatible with ErrorX as currently implemented. However, I plan to add support for alternate sequencing platforms, particularly long-read sequencing formats such as PacBio and Oxford Nanopore sequencing (Amarasinghe *et al.*, 2020), in the future. In the same vein, Sanger sequencing is also not currently supported, as the difference in sequencing chemistry and quality scores make it difficult to translate the results. Incorporating error correction for Sanger sequencing data is another future direction of ErrorX.

Currently UMI-based error correction methods have been used with great success in RepSeq experiments (Shugay *et al.*, 2014; Friedensohn *et al.*, 2018; Briney *et al.*, 2016; Khan *et al.*, 2016; Turchaninova *et al.*, 2016; Cole *et al.*, 2016). Incorporating a molecular barcode remains the gold standard for error correction as compared to bioinformatic approaches. However, UMI-based error correction faces many limitations that prevent its wide adoption, such as complex library preparation protocols, UMI collisions, loss of singleton sequences when samples are not sequenced to extraordinary depth, and lack of applicability to legacy data. Given these limitations, ErrorX can be used as a complementary approach to UMI barcoding to provide more confidence in the accuracy of sequences.

## Methods

### Datasets

Sequences were downloaded from the NCBI Sequence Read Archive (Leinonen *et al.*, 2011). Run identifiers were as follows: human BCR used in training data, SRR10413255, SRR10413256, SRR10413257, SRR10413258 (Sevy *et al.*, 2019); human BCR used in validation data, SRR10413259 (Sevy *et al.*, 2019); mouse BCR used in training data, SRR3174991, SRR3174992, SRR3175018, SRR3175020 (Khan *et al.*, 2016); mouse BCR used in validation data, SRR3175021 (Khan *et al.*, 2016); human TCR, SRR5676636 (Oakes *et al.*, 2017). Datasets were downloaded as separated paired end reads. Special care had to be taken during the paired end assembly process, since Phred quality scores are an input to the ErrorX neural network, and assembly softwares differ in their algorithms for calculating quality scores post-assembly. Therefore, to ensure that ErrorX would be robust to different paired end assemblers, reads were assembled using both PANDASEQ (Masella *et al.*, 2012) and USEARCH (Edgar, 2010) software packages. ErrorX was also tested on unassembled reads, with no change in performance.

### Data processing

Data was first assigned a germline sequence using IgBlast version 1.10 (Ye *et al.*, 2013). Sequences were discarded if the E value for V gene assignment was > 10^−4^. Sequences were kept in the dataset if the D and/or J gene assignments were poor (E value cutoffs were 10^−2^ and 10^−1^, respectively), but the putative D and/or J gene assignments were disregarded. As the public datasets used have incorporated barcodes, it was possible to determine precisely which positions were erroneous based on a consensus sequence. Reads were clustered based on a unique identifier consisting of the UMI barcode and V and J gene assignments to avoid barcode collisions. Clusters were then thrown out if they contained < 5 members to prevent an ambiguous consensus sequence. A consensus nucleotide sequence was then generated for each cluster and duplicate occurrences of the same consensus sequence were removed. This step was critical to ensure that the neural network would not overfit on the sequence of the most dominant clones. Error correction is a highly class-imbalanced problem, with roughly 0.5% positive cases. To enrich for positive cases, only non-germline positions were included in the training and validation sets, on the rationale that almost all positive cases were also non-germline. This filtering step enriched the proportion of positive cases to 3% total. After these processing steps a total of 2.4 × 10^6^ data points were included in the training set and 6.4 × 10^5^ data points in the validation set.

### Feature calculation

For each position in the nucleotide sequence, a sliding window of nine surrounding nucleotides (four preceding and four following the position of interest) was extracted and converted to numeric features using one-hot encoding. A sequence window based on the inferred germline sequence was also extracted, one-hot encoded and added to the feature vector. Additional features used as input were GC content, somatic hypermutation, and Phred quality scores, calculated both globally and within the sequence window. Phred quality score (*Q*) is a measure of the probability (*p*) of an erroneous base call by the sequencer, defined as: *Q* = −10log_10_*p*. A total of 124 features were used. Features were normalized between 0 and 1 before training.

### Training strategy

Data stratification into training and validation sets was performed using a “leave-one-donor-out” cross-validation strategy to provide the most rigorous testing possible. It has been observed that a traditional k-fold cross-validation strategy can be prone to overfitting when dealing with biological data, caused by batch effects between samples and multiple occurrences of the same sequence within a dataset (Leek *et al.*, 2010). Rather than pooling all sequencing across donors, then splitting into training/validation sets, the sequences comprising the training and validation sets were derived from separate donors, eliminating the possibility of cross-contamination. In addition, the human BCR, human TCR, and mouse BCR datasets were gathered by separate labs at different institutions - by including data gathered under different experimental conditions, the possibility that batch effects contribute to the performance of the ErrorX network was eliminated.

### Neural network architecture and training

After optimization of network architecture and type, the optimal network was determined to be a fully connected, feed forward multi-layer perceptron consisting of 3 hidden layers of 256, 128, and 64 nodes, respectively. Training was performed using a dropout layer between each hidden layer with a rate of 10%. In addition a kernel l2 regularizer was applied with a value of 10^−4^ to all hidden layers. ReLU activation functions were used for all layers except for the output neuron, which used a sigmoid activation function. A binary crossentropy loss function was used and optimized by a Stochastic Gradient Descent optimizer with a learning rate of 0.01. Training was performed using the Keras package with Theano backend in Python version 2.7 (The Theano Development Team *et al.*, 2016). The final network was trained on an r5.2xlarge AWS EC2 instance. Runtime calculations were performed on a MacBook Pro running OSX High Sierra 10.13.6, on an Intel i7 processing chip with multithreading enabled over 8 threads and 16 GB memory. ROC AUC and average precision score for the final predictions were calculated using the scikit-learn package (Pedregosa *et al.*).

### Clonal lineage reduction

A lineage of 20 sequences sharing a UMI barcode were identified from a mouse BCR sequencing dataset (Khan *et al.*, 2016; SRR3175021) that was part of the independent validation set during training. After germline gene assignment the nucleotide sequences were deduplicated to reduce the lineage to eight members. A phylogenetic tree was generated using the Interactive Tree of Life software (Letunic and Bork, 2019). After error correction by ErrorX the sequences were deduplicated again, ignoring N nucleotides (predicted sites of error).

### Software application

ErrorX is available as both a command-line application and a graphical user interface (GUI), called ErrorX Viewer. ErrorX is written in C++ and compiled using clang version 10.0.1. Unit tests covering all major functionality were written using the cxxtest package (https://cxxtest.com/). ErrorX Viewer is written in C++ and uses the Qt framework (https://www.qt.io/) version 5.10.1 for the interface. Both the command-line and GUI applications were compiled and tested on Mac OSX, Windows 7, Windows 10, and Debian-based Linux distributions for compatibility. Pre-compiled binaries are available at https://endeavorbio.com/downloads/, or from Github at https://github.com/EndeavorBio/ErrorX/releases for the command-line application and https://github.com/EndeavorBio/ErrorX-Viewer/releases for the GUI.

